# CCN2 activates ERK-signaling via integrin αv and enhances the interaction of ERK and DUSP6 in lymphatic endothelial cells

**DOI:** 10.1101/2021.06.18.449024

**Authors:** Shiho Hashiguchi, Tomoko Tanaka, Ryosuke Mano, Seiji Kondo, Shohta Kodama

## Abstract

Cellular communication network factor 2 (CCN2, also known as CTGF), is a modular and matricellular protein and a well-known angiogenic factor in physiological and pathological angiogenesis. However, its roles in lymphangiogenesis and intracellular signaling in lymphatic endothelial cells (LECs) remain unclear. Here, we investigated CCN2 signaling in LECs and its effects on lymphangiogenesis. In primary cultured LECs, gene expressions of lymphatic endothelial markers *lymphatic vessel endothelial hyaluronan receptor 1* (*Lyve1*), *Podoplanin* and *prospero homeobox 1* (*Prox1*) and lymphangiogenic factors *vascular endothelial cell growth factor c* (*Vegfc*), *vascular endothelial cell growth factor d* (*Vegfd*) and *fms-related tyrosine kinase 4* (*Flt4*, also known as *Vegfr3*) were upregulated by CCN2. Subsequently, we found that CCN2 induced phospho-ERK and that was decreased by suppression of integrin αv. CCN2 slightly decreased the growth of LECs due to enhancement of the interaction of ERK and dual specific protein phosphatase 6 (DUSP6), and knockdown of DUSP6 increased CCN2-induced phospho-ERK levels. In *in vivo* Matrigel plug assays, the number of Podoplanin-positive vessels was increased by exogenous CCN2, and phospho-ERK-positive LEC and DUSP6-positive LEC were detected in CCN2 plugs. These results suggest that CCN2-related lymphangiogenesis is regulated by DUSP6, which enables negative modulation of ERK-signaling.

## Introduction

Cellular communication network factor 2 (CCN2), also known as connective tissue growth factor (CTGF), was originally isolated as a platelet-derived growth factor (PDGF)-like factor in HUVEC (1). CCN2 is a member of the CCN family of modular matricellular proteins, which contains six family members (CCN1–6) (2). CCN family proteins contains four structural domains: the IGF binding domain, the von Willebrand factor C repeat domain, the thrombospondin type 1 (TSP) domain and the cysteine knot C-terminal domain (3). CCN2 is a multifunctional molecule, with roles in development, tumorigenesis, fibrotic disease and wound healing (4). Of note, CCN2 has been also considered to be one of the most important regulators of both physiologic and pathologic angiogenesis (5-7). CCN2/CTGF is highly expressed in vasculature, including in the coronary vessels of the heart and vasculature of the kidneys, lung and liver (8-10), and in certain tumors to promote tumor angiogenesis (5,11).

With regards to the CCN2 signaling cascades, CCN2 activates MAPKs such as p38 MAPK, ERK, and JNK in HUVEC and chondrocytes, either directly via cell-surface receptors or indirectly via other extracellular cofactors. (12,13) Integrins are cell surface adhesion receptors for extracellular matrix proteins and immunoglobulin superfamily proteins, and conformation change of the extracellular domain triggers activation of intracellular signaling pathways, including ERK and MAPK signaling(14-17). The TSP domain in CCN2 interacts with integrin α6β1(18), and the C-terminal motif of CCN2 interacts with integrins αvβ33, α5β31 and α6β31 (19-24). In vascular endothelial cells, CCN2/CTGF functions through direct binding to integrin αvβ33 (19,25) to promote proliferation and induce chemotaxis and formation of tubules (26,27).

The lymphatic vascular system has essential roles in the regulation of tissue pressure, immune surveillance and absorption of dietary fat in the intestine (28). Lymphangiogenesis is necessary not only for the development of lymphatic system during embryogenesis, but also for tissue repair and inflammatory reactions in most organs (29). Several angiogenic factors are involved in growth of lymphatic vessels, such as the Tie/angiopoietin system, neuropilin-2 and integrin α9, which regulate lymphangiogenesis (30-32). VEGF-C-VEGFR-3 signaling is also crucial for the development of lymphatic vessels (33-36). Impairment of lymphatic function results in various diseases that are characterized by the inadequate transport of interstitial fluid, edema, impaired immunity, and fibrosis (37,38). Abnormal proliferation of lymphatic endothelial cells (LECs) causes lymphangiomas, lymphangiosarcomas and Kaposi’s sarcoma (39,40).

Previous studies in renal, peritoneal dialysis-related and liver fibrosis mouse models showed that CTGF enhanced lymphangiogenesis (41-44). Furthermore, CTGF is involved in fibrosis-associated renal lymphangiogenesis through its interaction with VEGF-C (41,45). These reports show that CCN2 is closely associated with lymphangiogenesis in fibrous diseases. However, CCN2 signaling in LECs in lymphangiogenic signaling has not been elucidated. In this study, we investigated the downstream signaling of CCN2 in primary cultured LECs.

## Results

### CCN2 increases the gene expression levels of LEC markers and *Vegfc, Vegfd and Flt4* (*Vegfr3*) in LECs

To investigate the roles of CCN2 in lymphangiogenic signaling, we examined the gene expression levels of LEC markers, *lymphatic vessel endothelial hyaluronan receptor 1* (*Lyve1*), *Podoplanin* (*Pdpn*) and *prospero homeobox 1* (*Prox1*), and lymphangiogenic factors, *vascular endothelial cell growth factor c* (*Vegfc*), *vascular endothelial cell growth factor d* (*Vegfd*), *fms-related tyrosine kinase 4* (*Flt4*, also referred to as *Vegfr3*) and *kinase insert domain receptor* (*Kdr* also referred to as *Vegfr2*), in primary cultured LECs treated with CCN2. LECs were stimulated with 0, 12.5, 25, 50, 100 or 200 ng/mL of CCN2 for 3 h, and the expression levels of the LEC markers *Lyve1, Pdpn* and *Prox1* were analyzed by quantitative RT-PCR. The expression levels of *Lyve1, Pdpn* and *Prox1* were increased by CCN2 in a dose-dependent manner up to 100 ng/mL; at 200 ng/mL, their expression levels dramatically decreased compared with levels at 100 ng/mL (Fig. 1A). We further found that CCN2 increased the expression levels of the lymphangiogenic factors *Vegfc, Vegfd* and *Flt4* but did not affect *Kdr* expression (Fig. 1B). Phosphorylation of the receptor tyrosine kinases (RTKs) VEGFR2 and VEGFR3 were investigated using a phospho-RTK array. Our results showed that VEGFR2 and VEGFR3 phosphorylations were unchanged by CCN2 (Fig. S1). In contrast, phosphorylation levels of the RTKs EGFR and EphB4 were increased by CCN2, while phosphorylation of ErbB4 and PDGFRα were decreased compared with controls (p<0.001).

**Figure 1.**
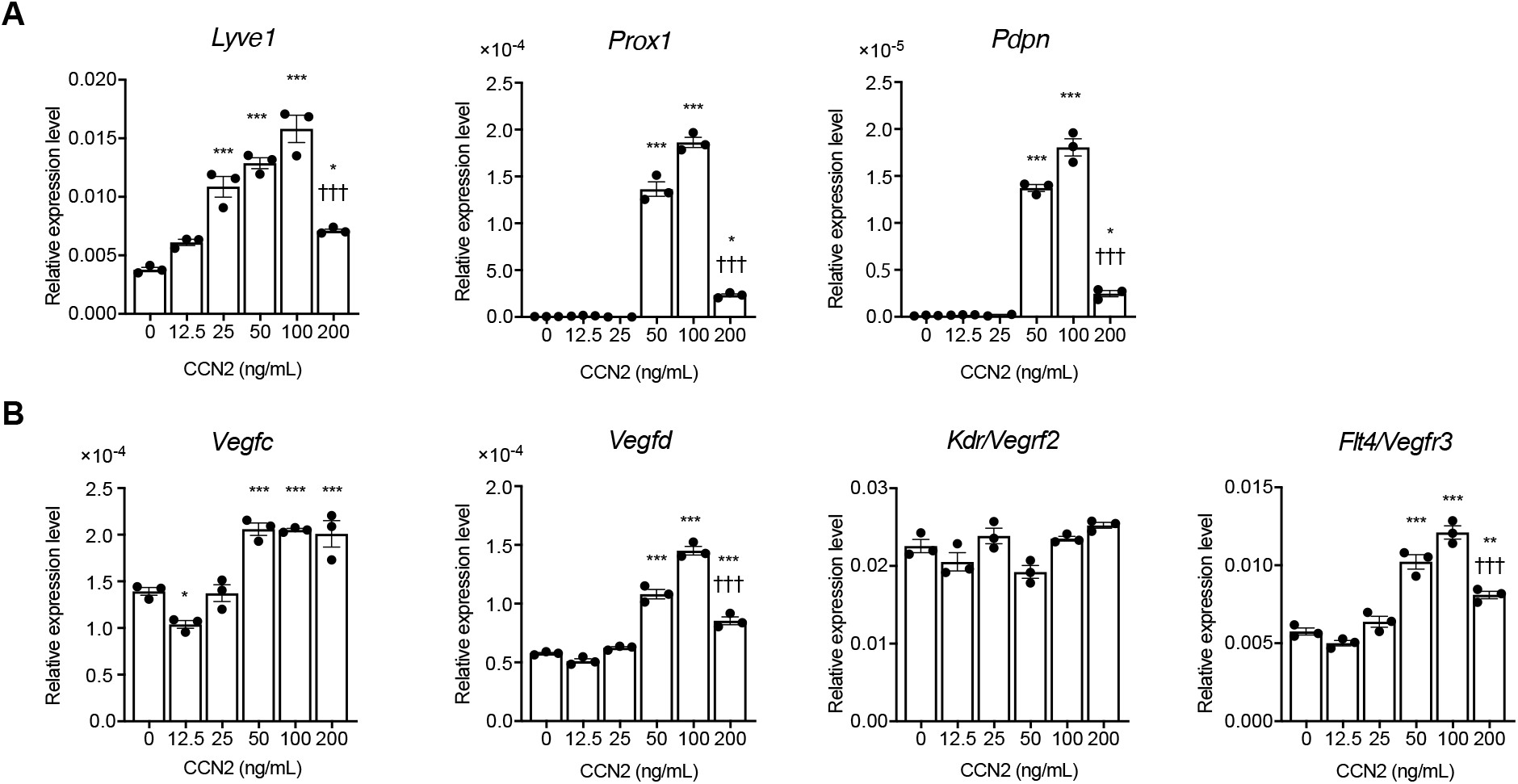
CCN2 upregulates lymphangiogenesis markers and *Vegfc, Vegfd* and *Flt4* (*Verfr3*) in lymphatic endothelial cells (LECs). Mouse primary cultured LECs were treated with 0, 12.5, 25, 50, 100 or 200 ng/mL CCN2 for 3 h. (A) mRNA expression levels of *Lyve1, Podoplanin* and *Prox1* were analyzed by quantitative RT-PCR. (B) *Vegfc, Vegfd, Kdr* (*Vegfr2*) and *Flt4* (*Vegfr3*) mRNA levels were analyzed by quantitative RT-PCR. *Actb* was used as an internal control, and gene expression levels were expressed relative to *Actb* mRNA levels. One-way ANOVA, ^*^p < 0.05, ^**^p < 0.01, ^***^p < 0.01 vs. 0 ng/mL Ccn2. ^†††^p < 0.01 vs. 100 ng/mL CCN2.

### CCN2 enhances the phosphorylation of ERK via integrin αv in LECs

To determine the intracellular signaling downstream of CCN2, we next analyzed the expression of various phosphorylated proteins that are known to be activated by CCN2 stimulation (27). While the phosphorylation levels of p38 MAPK, JNK, AKT, NFκB and Smad2, -3, and -1/5 were unchanged in response to CCN2 (Fig. S2A–C), phospho-ERK was significantly increased in response to CCN2 (p<0.001) (Fig. 2A, 2B). Total ERK protein levels were not affected by CCN2 treatment (Fig. 2C).

**Figure 2.**
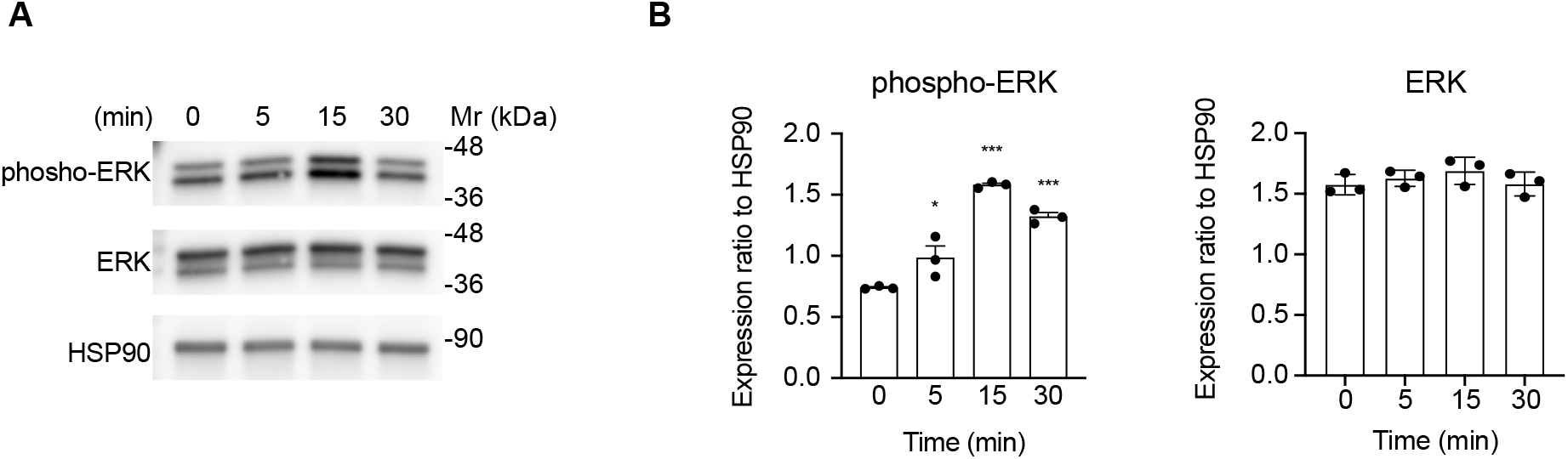
Phosphorylation of ERK is increased by CCN2. (A) LECs were stimulated with 100 ng/mL CCN2 for 0, 5, 15 or 30 min, and phosphorylated ERK, total ERK and HSP90 were detected by Western blotting. (B) Quantification of protein bands from (A); phospho-ERK and total ERK levels were normalized with HSP90. One-way ANOVA, ^*^p < 0.05, ^**^p < 0.01, ^***^p < 0.001 vs. 0 min.

Although Vegfrs are upstream of ERK (46,47), our RTK assay results showed that VEGFR2 and VEGFR3 were not activated by CCN2 in LECs. Previous studies showed that CCN2 promotes vascular endothelial cell growth, migration, adhesion, and survival partly through its interactions with integrins (25). Therefore, we next investigated whether CCN2-mediated effects on ERK may involve its interaction with integrins. We first examined the expression levels of *Itgav, Itga9, Itga2b, Itgb1, Itgb2* and *Itgb3* mRNA in primary LECs treated with CCN2. *Itgav, Itga9 and Itgb*1 were predominantly expressed in LECs. Furthermore, the expression levels of these mRNAs were increased by CCN2 stimulation (Fig. 3A). *Itga2b, Itgb2* and *Itgb3* were not induced by CCN2. Knockdown of integrin αv by siRNA partially decreased phosphorylated ERK induced by CCN2 compared with non-target siRNA (p=0.0040) (Fig. 3B), which suggests that CCN2 increases phospho-ERK via integrin αv in LECs. In contrast, knockdown of integrin α9 resulted in a slight increase of CCN2-induced phospho-ERK compared with levels in cells transfected with non-target siRNA (Fig. 3C). Interestingly, phospho-ERK was increased by suppression of integrin β1 (p=0.05) but was not altered by CCN2 stimulation in integrin β1– suppressed LECs (Fig. 3D). *Itgb2* and *Itgb3* mRNA levels were not changed by suppression of integrin β1, which suggests that integrins β2 and β3 may not compensate for the decrease in integrin β1 (Fig. S3).

**Figure 3.**
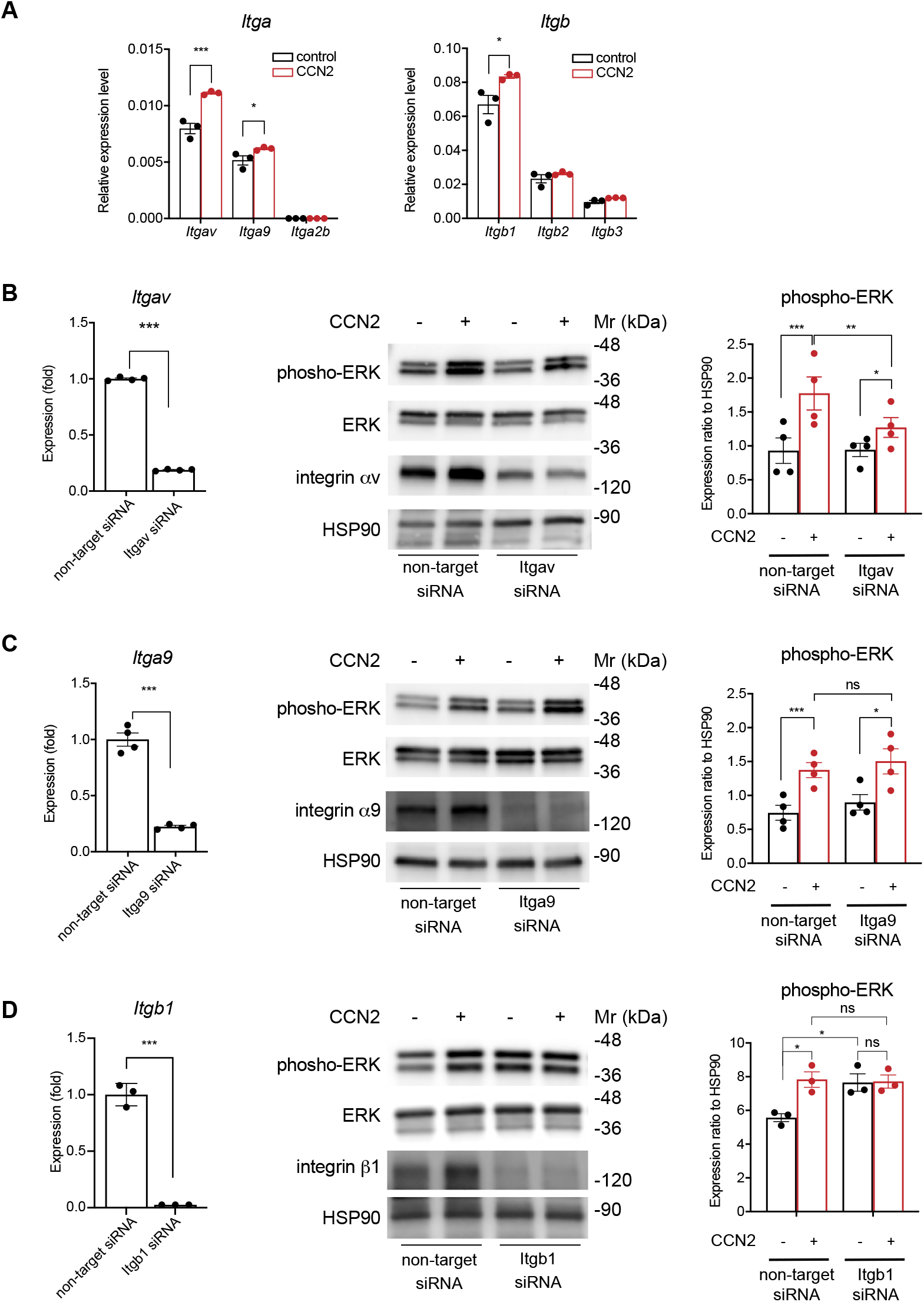
CCN2 induces phospho-ERK levels via integrin αv. (A) LECs were treated with or without 100 ng/mL CCN2 and mRNA expression levels of integrin family members were evaluated by quantitative RT-PCR. Gene expressions were normalized to levels of *Actb* mRNA. *Itgav, Itga9* and *Itga2b* mRNA levels and *Itgb1, Itgb2* and *Itgb*3 results are shown in left and right panels, respectively. Two-way ANOVA, ^*^*p* < 0.05, ^***^*p* < 0.001. Suppression of integrin αv (B), α9 (C) and β1 (D) by siRNA. mRNA expression levels of integrin (left), Student t-test, ^***^p < 0.001 vs. non-target siRNA. Western blot in LEC transfected with siRNA and treated with CCN2 (middle). Phospho-ERK levels in the middle were quantified and normalized with HSP90 expression levels (right), two-way ANOVA, ^*^p < 0.05, ^**^p < 0.01, ^***^p < 0.001. ns; not significant.

### CCN2 promotes the binding of ERK and DUSP6 in LECs

CCN2 promotes cell proliferation in vascular endothelium (26,48-50). Therefore, we next analyzed the effects of CCN2 on the cell growth of LECs. Treatment of LECs with 10, 50 and 100 ng/mL of CCN2 for 24 h suppressed cell growth compared with cells at 0 h (p = 0.22, 0.92 and > 0.99, respectively) (Fig. 4A). In addition, treatment with 50 and 100 ng/mL of CCN2 suppressed cell growth at 24 h compared with vehicle-treated LECs (p = 0.003 and 0.02). However, no changes in proliferation were observed at 48 and 72 h of treatment. These results indicate CCN2 has a weak growth inhibitory effect on LECs.

**Figure 4.**
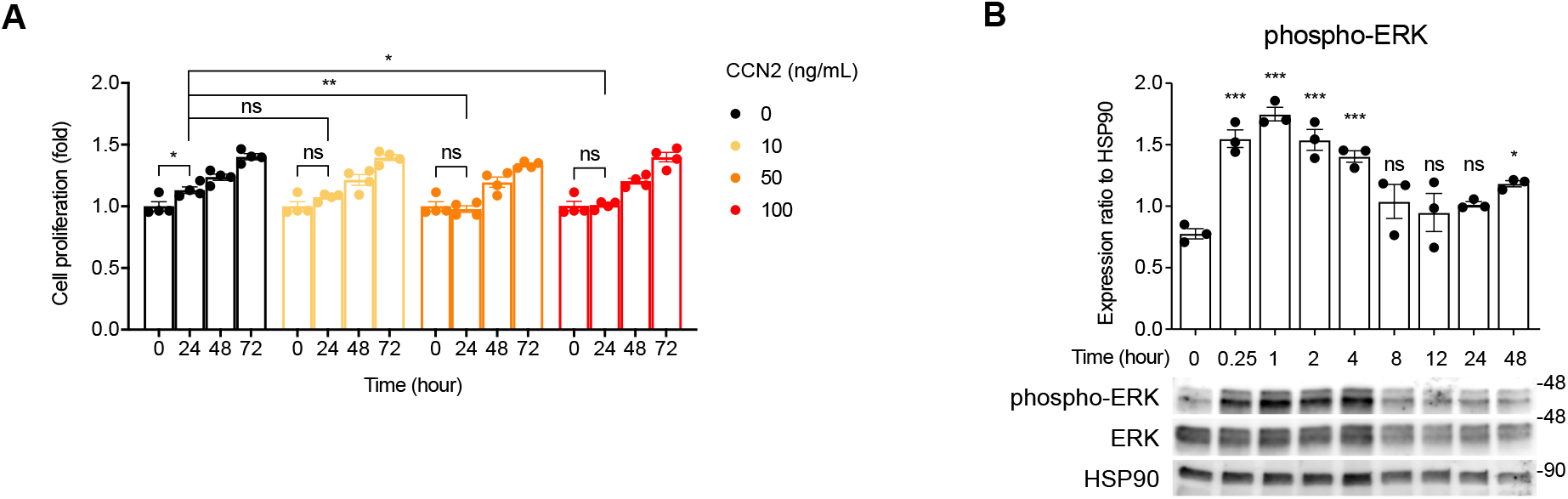
Weak inhibitory effects of CCN2 on the proliferation of LECs and transient phosphorylation of ERK induced by CCN2. (A) LECs were cultured in medium containing 0, 10, 50 or 100 ng/mL of CCN2 for 24, 48 and 72 h. Cell proliferation was analyzed by crystal violet staining. Cell proliferation was expressed as the ratio to the value at 0 h. Two-way ANOVA, ^*^p < 0.05, ^**^p < 0.01. ns; not significant. (B) LECs were treated with 100 ng/mL CCN2 for 0, 0.25, 1, 2, 4, 8, 12, 24 or 48 h, and phospho-ERK, ERK and HSP90 were detected by Western blot. Phospho-ERK levels at each time point were normalized to those of HSP90. One-way ANOVA, ^*^p < 0.05, ^***^p < 0.001 vs. 0 h. ns; not significant vs. 0 h.

Time-course experiments showed that phospho-ERK was significantly increased in LECs at 15 min after treatment with CCN2 and continued to increase up to 4h, followed by a decrease at 8 h, and increase after 48 h (Fig. 4B). The phosphorylation of ERK induced by CCN2 was non-persistent and transient, suggesting the presence of a concomitant mechanism for suppressing ERK.

The inactivation of MAPKs by dephosphorylation of threonine and/or tyrosine residues of the T-X-Y motif within the kinase activation loop is mediated by serine/threonine phosphatases, tyrosine phosphatase, and dual-specificity phosphatases (DUSPs) (51). The DUSP family phosphatases are the largest group of protein phosphatases that specifically regulate MAPK activity in mammalian cells (51). Quantitative RT-PCR revealed that *Dusp1, -2, -4, -5, -6, - 7, -8, -9* and -*10* are expressed in LECs; *Dusp6* was predominant, but *Dusp16* was undetected (Fig. 5A).

**Figure 5.**
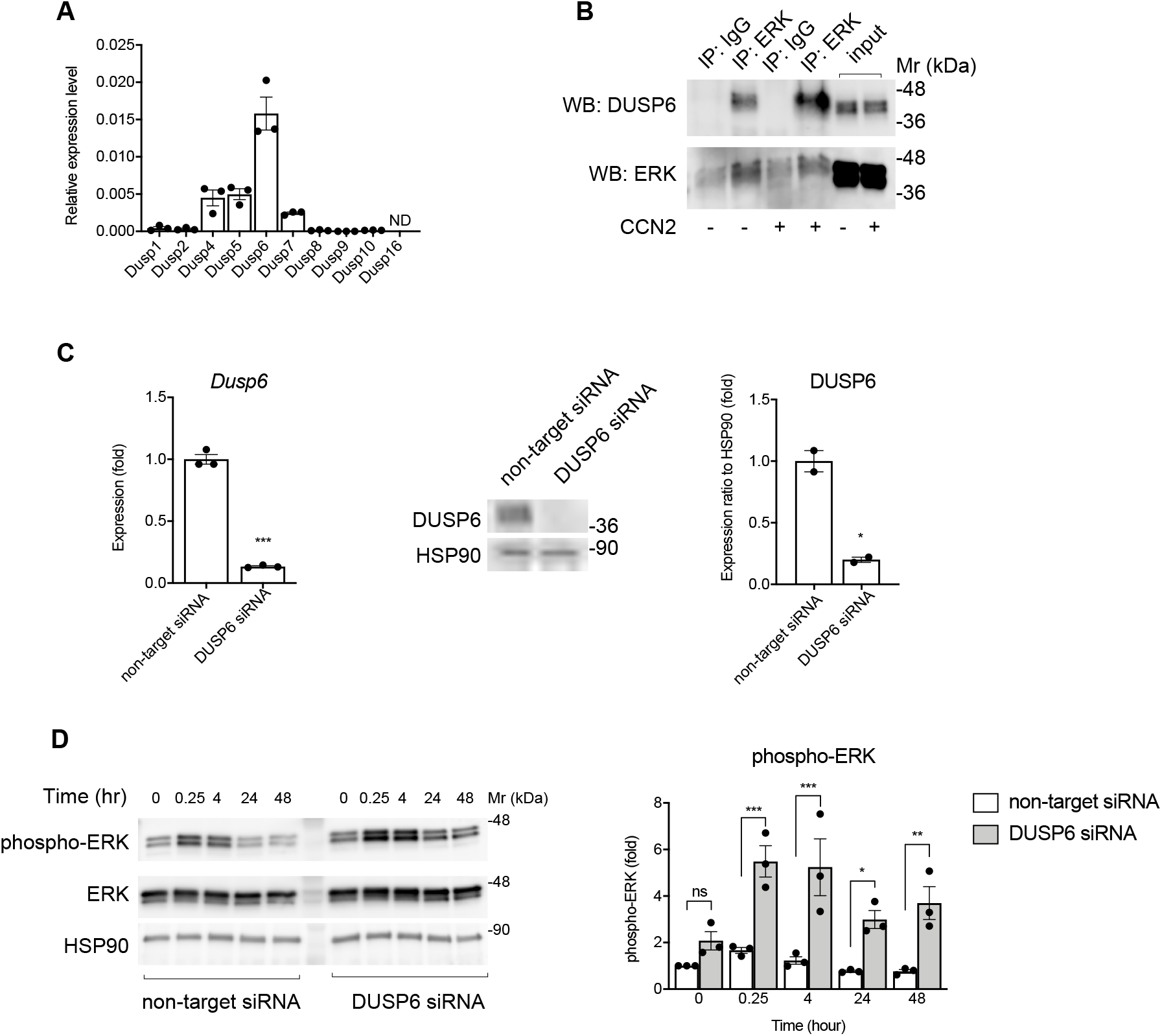
DUSP6 interacts with ERK and suppression of DUSP6 increases phospho-ERK levels in LECs. (A) Expression levels of *Dusp1, -2, -4, -5, -6, -7, -8, -10* and *-16* in LECs. ND, not detected. (B) Cell lysates from LECs stimulated with 100 ng/mL of CCN2 for 5 min or control cells were immunoprecipitated with anti-ERK antibody (ERK) or normal rabbit IgG (IgG) and analyzed by Western blot with antibodies against DUSP6 or ERK. (C) Suppression of DUSP6 by siRNA. mRNA expression levels (left) and protein levels (middle) in LECs transfected with non-target siRNA or DUSP6 siRNA. Quantification of protein bands from middle; DUSP6 level was normalized with HSP90. (D) Phospho-ERK levels in LECs transfected with non-target siRNA or DUSP6 siRNA, and treated with 100 ng/mL CCN2 for 0, 0.25, 4, 24 and 48 h. phospho-ERK levels were quantified and normalized with HSP90 levels. Two-way ANOVA, ^*^p < 0.05, ^**^p < 0.01, ^***^p < 0.001, vs. non-target siRNA. ns, not significant.

Binding of activated ERK to the kinase interaction motif of DUSP6 results in a conformational change, prompting phosphatase activation of the DUSP6 catalytic domain, leading to dephosphorylation of ERK (52). To investigate whether ERK and DUSP6 directly interact, we performed immunoprecipitation assays. The results showed that DUSP6 co-precipitated with ERK, and their interaction was enhanced by CCN2 (Fig. 5B). In addition, in LECs transfected with siRNA targeting DUSP6 (Fig. 5C), phospho-ERK levels were increased compared with levels in cells transfected with non-target siRNA at 15 min and 4, 24 and 48 h after CCN2 treatment. (p<0.001 for 15 m, 4 h,; p=0.04 for 24 h,; p=0.004 for 48 h) (Fig. 5D). These results indicate that while CCN2 enhances the phosphorylation of ERK, CCN2 also enhances the binding between ERK and DUSP6, and DUSP6 induces dephosphorylation of ERK in LECs.

### CCN2 promotes lymphangiogenesis in *in vivo* Matrigel plug assay

To investigate the effects of CCN2 on lymphangiogenesis *in vivo*, recombinant CCN2 was mixed with Matrigel and subcutaneously injected onto the backs of mice. At 7 days after injection, blood vessels were macroscopically observed on the surface of plugs in the CCN2 group mice, but not in the control group (Fig. 6A). Histological analyses revealed that the cells in the CCN2 gel were mostly Podoplanin-positive, and tubular structures were observed (Fig. 6B and 6C). The number of Podoplanin-positive vessels and percentage of Podoplanin-positive areas were increased in CCN2-supplemented gels compared with controls, which suggests that CCN2 positively modulated lymphangiogenesis. CCN2 also increased the number of CD31-positive cavities and β3-Tubulin-positive peripheral nerves in the CCN2 plugs (Fig. S4). Immunostaining revealed that total ERK was similarly expressed in both the control gel and the CCN2 gel; however, while phosphorylated ERK was barely present in the control gel, it was detected in the CCN2 gel and localized in the nucleus and cytosol of Podoplanin-positive cells (Fig. 7A and 7C). DUSP6 was also rarely detected in the control gel, but it was detected in the cytosol of Podoplanin-positive cells in the CCN2 gel (Fig. 7B). Phosphorylated ERK and DUSP6 expression in Podoplanin-positive cells were significantly increased in the CCN2 gel compared with the control gel (p < 0.05) (Fig. 7A and 7B).

**Figure 6.**
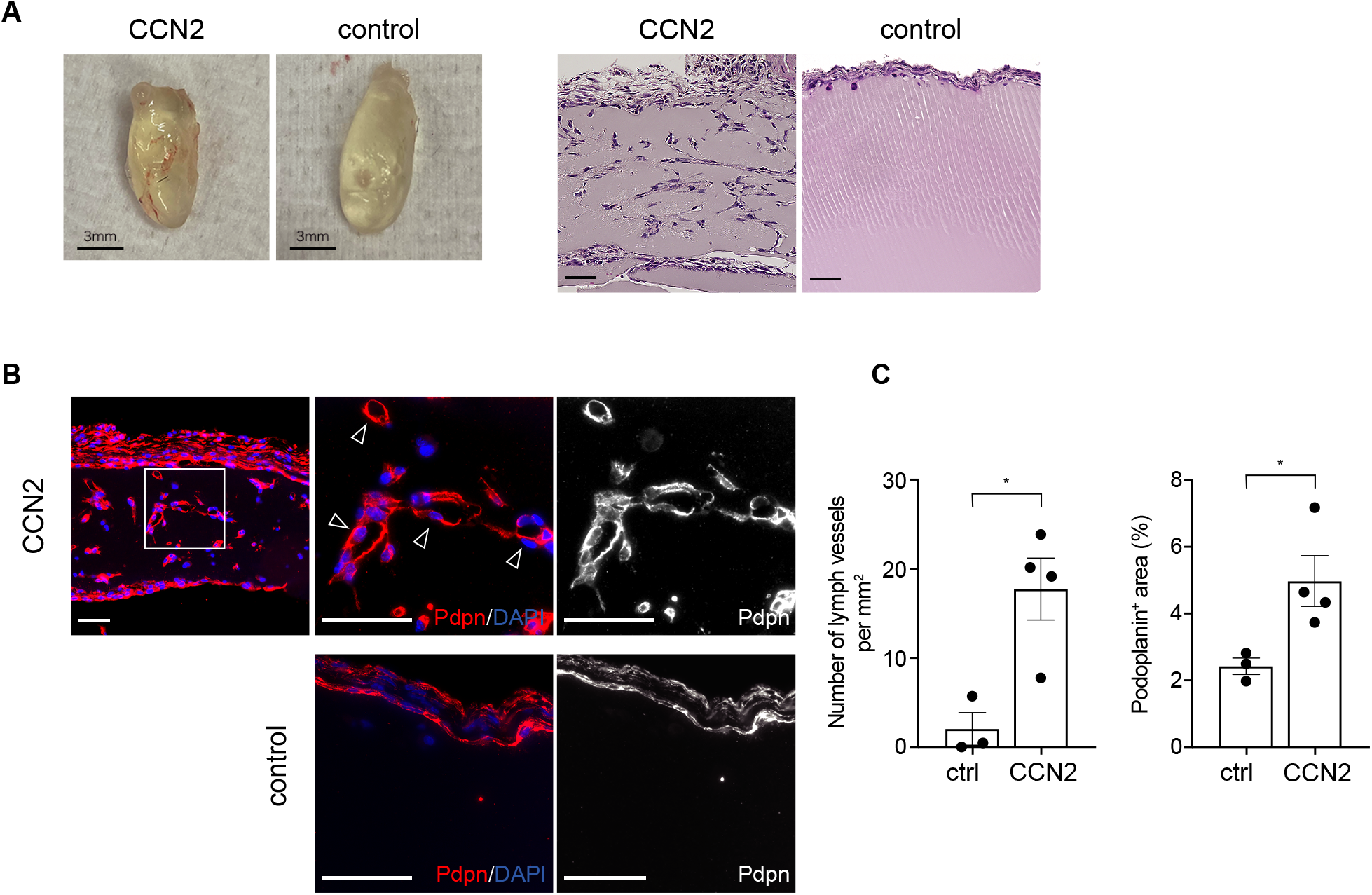
CCN2 enhances lymphangiogenesis in *in vivo* Matrigel plug assay. Matrigel and recombinant CCN2 were mixed and injected subcutaneously back of the mice. Control mice were injected with Matrigel and PBS mixture. (A) Images and HE-stained sections of Matrigel removed 7 days after injection. Scale bars: 3 mm (images) and 50 *µ*m (HE-stained sections). (B) Immunostaining of Podoplanin. Podoplanin-positive cells on the surface side of control gel and inside of the CCN2 gel. Arrowheads indicate Podoplanin-positive vessels in the CCN2 gel. Bars: 50 *µ*m. (C) Number of Podoplanin-positive vessels per mm^2^ and the ratio of the Podoplanin-positive area to the whole area. Student t-test, *p < 0.05.

**Figure 7.**
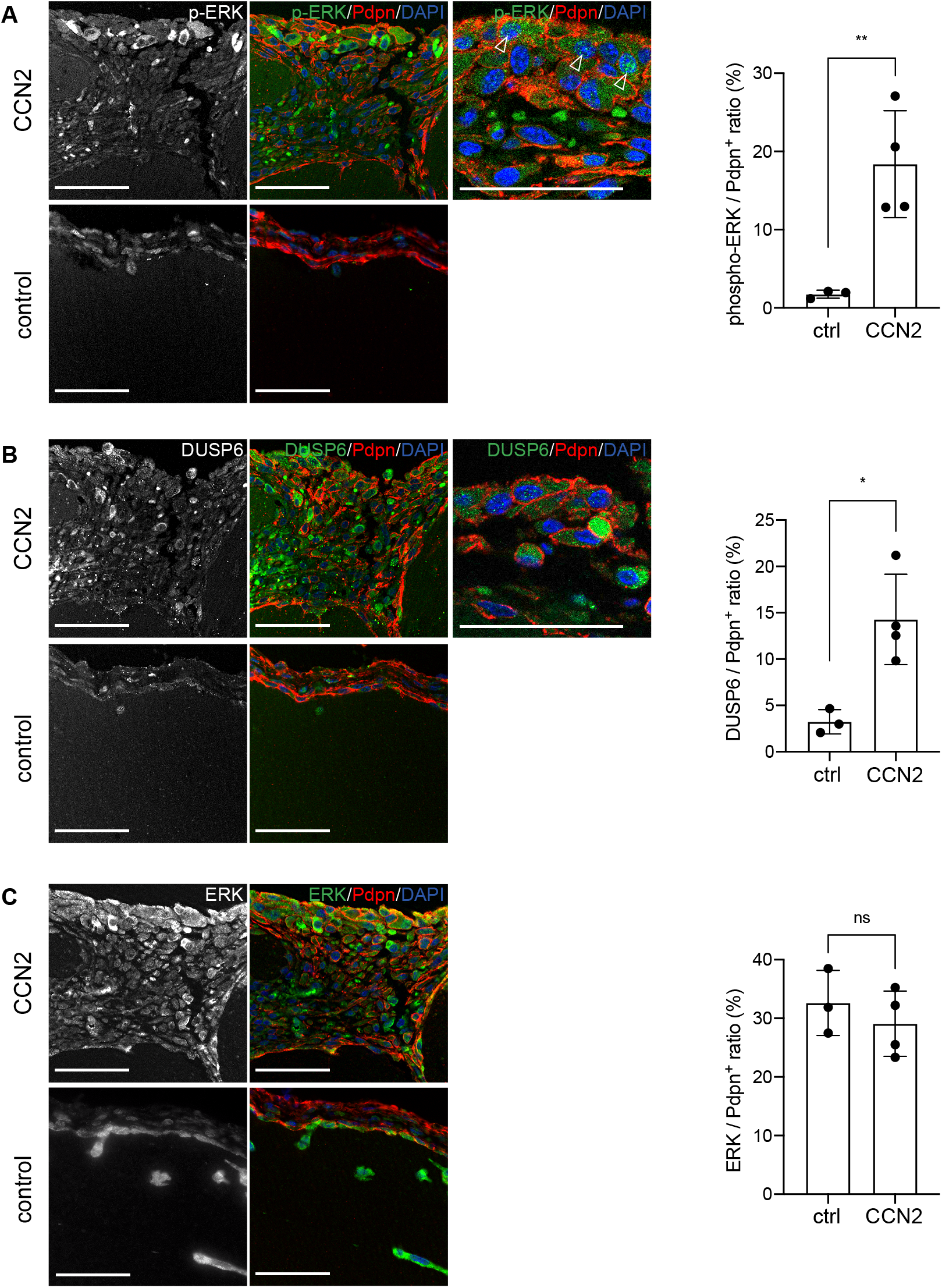
Enhanced expression of phospho-ERK and DUSP6 in the CCN2 matrigel. Expression of phospho-ERK, DUSP6 and ERK in Matrigel plugs detected by immunostaining (left) and quantification of the ratio of the phospho-ERK, DUSP6 or ERK-positive area to the Podoplanin-positive area for each gel (right). (A) Phospho-ERK-positive cells were localized on the surface side of the CCN2 gel. Arrowheads indicate Podoplanin-positive/phospho-ERK-positive cells. DUSP6-positive cells (B) and ERK-positive cells (C) were detected in both the CCN2 gel and control gel. Scale bars: 50 *µ*m. Student t-test, *p < 0.05, ^**^p < 0.01. ns; not significant.

## Discussion

The molecular mechanism underlying the cell proliferation of LECs involves the activation of ERK signaling stimulated by autocrine or paracrine VEGF-C and VEGFR3 signaling (53). We demonstrated that CCN2 upregulates *Vegfc, Vegfd* and *Flt4* and increases the phosphorylation of ERK in primary cultured LECs. However, the phosphorylation of VEGFR2 (KDR) and VEGFR3 (FLT4) was not increased by CCN2, which suggests that CCN2 and endogenous VEGF-C secreted from LECs do not activate VEGF receptor signaling upstream of ERK in primary cultured LECs.

Integrins play key roles in endothelial cell migration and survival during angiogenesis and lymphangiogenesis. A previous study showed that integrin αvβ3 enhanced angiogenesis induced by basic FGF and TNFα, and integrin αvβ5 enhanced VEGF-induced angiogenesis and tumor metastasis (54). Integrin α9β1 is required for proper development of the lymphatic system (54). In agreement with these studies, integrin αv, α9 and β1 were detected in LECs, and CCN2-induced phospho-ERK was reduced by suppression of integrin αv, whereas suppression of integrin α9 did not decrease phosphorylated ERK. These results indicate that CCN2-induced phospho-ERK is partially mediated by integrin αv in LECs. Recently, Kumaravel et al. showed that integrin β1 was important in lymphangiogenesis (55). However, phospho-ERK was increased by suppression of integrin β1 without CCN2 stimulation. The loss of β1 was not compensated for by increases in integrins β2 and β3. In LECs, integrin α5 is increased by suppression of integrin β1 (55). We did not investigate the expression level of *Itga5* mRNA because the interaction between CCN2 and integrin α5 has not been reported in LECs. The increase in phospho-ERK without CCN2 stimulation might be due to integrin α5 supplementation. However, phospho-ERK was not altered by CCN2 stimulation in integrin β1 suppression, suggesting that integrin β1 might be involved in CCN2 signaling in LECs.

Lymphangiogenesis in adults involves the cell proliferation, budding and migration of LECs. Our *in vitro* cell proliferation assay results showed that CCN2 did not increase the proliferation of LECs but rather weakly suppressed proliferation. In HUVECs, DUSP1 and DUSP5 regulate the MAPK signaling pathways that are activated by VEGF (56), and DUSP5 modulates ERK1/2 downstream of the Ang1/Tie2 receptors (57). We showed DUSP6 is highly expressed in LECs, and immunoprecipitation assays showed that CCN2 enhanced the interaction of ERK and DUSP6 in LECs. Moreover, suppression of DUSP6 by siRNA increased phospho-ERK levels compared with control cells, which suggests that DUSP6 functions as a negative modulator of ERK-signaling in LECs. In a previous study, Forkhead box C1 (FOXC1) and Forkhead box C2 (FOXC2) were identified as negative regulators of ERK (58). In mice, lymphatic endothelial cell-specific deletion of FOXC1, FOXC2, or both resulted in lymphatic hyperplasia via hyperactivation of ERK (58). Here we identified DUSP6 as a suppressor of ERK in LECs; however, the relationship between FOXC1, FOXC2 and DUSP6 has not yet been analyzed.

Our results showed that CCN2 enhanced lymphangiogenesis in *in vivo* Matrigel assays, similar to findings in fibrosis model studies (43). The expression of phosphorylated ERK and DUSP6 was similarly strong in Podoplanin-positive cells in the CCN2 gel, which suggests that DUSP6 might suppress the excessive activation of ERK, as shown *in vitro*. CCN2 also enhanced angiogenesis and peripheral nerve cell migration. Previous angiogenesis studies showed that CCN2 is sufficient to induce angiogenesis, directly or indirectly, under *in vitro* and *in vivo* experimental conditions (19,26,59). CTGF expression increases after central nervous system trauma in rodent models (60-62). In zebrafish spinal cord injury models, injury-induced ctgfa directs glial cell bridging and neuronal regeneration (63). Our results are consistent with these previous studies. The discrepancy observed between the *in vitro* cell proliferation and *in vivo* Matrigel assays may be due to effects of ECM and matrix metalloproteinase (MMP) in Matrigel. Since CCN2-signaling is mediated by ECM and activated through proteolytic processing by MMP-2,3 (6,64-67), lymphangiogenesis in *in vivo* Matrigel assays was promoted. The results between these assays may not be comparable, and our present results are not contradictory.

A recent study found significant expression of CTGF in the epithelium and connective tissue of oral submucous fibrosis, which was classified as a potentially malignant oral disease, as well as oral squamous cell carcinoma (OSCC) cases, increasing with disease progression, whereas healthy buccal mucosa had no CTGF expression (68). Moreover, MSC-derived CCN2 promoted tongue squamous cell carcinoma progression *in vitro* and *in vivo*. CCN2 is expressed at higher levels in tongue squamous cell carcinoma tissues than in paraneoplastic tissues, and its expression correlated with lymph node metastasis (69). A study of patient-derived tumor samples in the Cancer Genome Atlas database showed that *CCN2* expression was higher in head and neck cancers, and high expression of *CCN2* and *MMP3* was associated with worse prognosis in head and neck cancers (64). Lysine-specific demethylase 1 (LSD1) is a nuclear histone demethylase that functions as an epigenetic regulator to promote cancer initiation, progression, and relapse. LSD1 inhibitors markedly downregulated *CTGF* mRNA and protein expression and inhibited further xenograft growth in a tonsillar OSCC patient-derived xenograft mouse model (70).

We demonstrated that CCN2 activates ERK signaling via integrin αv and promotes lymphangiogenesis. In addition, CCN2 enhances the binding of DUSP6 to ERK, and DUSP6 suppresses excessive CCN2-induced phosphorylation of ERK. In OSCC, CCN2-signaling is associated with disease progression and lymph node metastasis. Our results suggest that integrin αv, ERK and DUSP6-mediated lymphangiogenesis may be therapeutic targets of OSCC.

## Experimental procedures

### Cell culture and proliferation assay

C57BL/6 mouse primary cultured LECs (CellBiologics, Chicago, IL, USA) were cultured in 0.1% gelatin-coated culture dishes at 37°C with 5% CO_2_. LECs were cultured using the EGM2-Endothelial cell growth medium-2 bullet kit (Lonza, Basel, Switzerland). Cells up to 6 passages were used in *in vitro* experiments. Cell proliferation was assessed by crystal violet staining method. Cells were plated in 24-well plates at 2 × 10^4^ cells per well and cultured overnight. The medium was changed to medium containing 0, 10, 50 or 100 ng/mL recombinant rat CCN2/CTGF protein, Carrier Free (R&D Systems, Minneapolis, MN, USA). At 0, 24, 48 and 72 h after treatment, cells were fixed with methanol, stained with 0.5% crystal violet (Merck, Darmstadt, Germany) in 25% methanol for 5 min, washed with water five times and dried. Absorbance at 590 nm was measured using the SPARK 10M plate reader (TECAN, Mannedorf, Switzerland). Cell proliferation was expressed as the ratio of the absorbance of each time point to the absorbance at 0 h.

### Quantitative RT-PCR

Total RNA was prepared from cells using the Purelink RNA purification Kit (Invitrogen, Carlsbad, CA, USA). cDNA was prepared from 1 μg of total RNA using the QuantiTect RT Kit (Qiagen, Venlo, the Netherlands). Quantitative RT-PCR was performed using TB Green Premix ExTaq II (Takarabio, Otsu, Japan) and a LightCycler96 (Roche Basel, Switzerland). Primers used for qPCR are listed in Table S1.

### Gene silencing by siRNA

Expression of integrins and DUSP6 was suppressed by siRNA. siRNAs were designed by an siRNA design site, siDirect ver.2 (http://sidirect2.rnai.jp/). Sense and complementary strand RNAs having a two-mer overhang were synthesized and annealed. The siRNA sequences for each mRNA target are listed in Table S2. LECs (1.5 × 10^5^) were transfected with 25 pmol of target siRNA or non-target siRNA (MISSION siRNA universal control#1, Merck) using Lipofectamine RNAiMAX (Thermo Fisher, Waltham, MA, USA) according to the manufacturer’s instructions.

### Western blot analysis

LECs were pre-cultured with FBS-free and supplement-free EGM2 medium (Lonza) for 24 h. Cells were then treated with CCN2-containing FBS-free and supplement-free EGM2 medium and cultured for the appropriate time. Cells were washed with PBS and lysed in RIPA Buffer (Merck) containing cOmplete protease inhibitor cocktail (Roche) and phosphatase inhibitor cocktail (Nacalai, Kyoto, Japan). Protein concentration was determined with the BCA protein assay kit (Thermo). Protein samples (5–20 *µ*g) were separated with SDS-PAGE and transferred to PVDF membranes (Invitrogen). The membrane was blocked in Blocking-one (Nacalai) for 30 min and then incubated with primary antibodies at 4°C overnight. The primary antibodies are listed in Table S3. After three washes with PBS containing 0.1% Tween-20, the membrane was incubated with HRP-conjugated anti-rabbit IgG or HRP-conjugated anti-rat IgG secondary antibody (Cell Signaling Technology, Danvers, MA, USA) for 2 h. After three washes with PBS containing 0.1% Tween-20, signals were visualized using ECL Prime (GE Healthcare, Chicago, IL, USA) and an image analyzer 680 (GE Healthcare). Signals were quantified using ImageQuant TL 8.1 software (GE Healthcare).

### Phospho-receptor tyrosine kinase array

LECs were pre-cultured with FBS-free and supplement-free basal medium for 24 h and then treated with medium containing 100 ng/mL of recombinant rat CTGF (R&D Systems) for 5 min. The phosphorylation levels of RTK were analyzed using a Proteome profiler mouse phospho-RTK array kit (R&D Systems) according to the manufacturer’s instructions. Briefly, 200 *µ*g of total protein was incubated with the receptor-spotted membrane at 4°C for 24 h. After washing, the membrane was incubated with anti-phospho tyrosine kinase antibody, and the phosphorylation levels of receptor tyrosine kinase were detected with the image analyzer 680 (GE Healthcare).

### Immunoprecipitation assay

Cells were lysed in Lysis Buffer (CST) containing cOmplete protease inhibitor cocktail and phosphatase inhibitor cocktail. Protein concentration was determined with the BCA protein assay kit (Thermo). Cell lysates containing 200 *µ*g of total protein were incubated with anti-ERK (CST) or anti-DUSP6 (Abcam, Cambridge, UK) antibody at 4°C overnight. Protein A magnetic beads (New England Biolabs, Ipswich, MA, USA) were then added to the cell lysate and antibody mixture, and the samples were mixed for 20 min. Magnetic beads were separated and washed with lysis buffer five times; the samples were denatured in 20 *µ*l of 3×SDS Sample Buffer and subjected to Western blotting.

### *In vivo* Matrigel plug assay

Male 8-week-old C57BL/6 mice were purchased from Charles River Japan. Mice were kept under specific pathogen-free conditions and used at 9 weeks of age. All experiments were performed in compliance with the relevant laws and institutional guidelines and were approved by the Animal Care and Use Committee of Fukuoka University (approval number: 2008024).

We mixed 450 *µ*L of growth factor reduced Matrigel (Corning, Corning, NY, USA) and 8.33 *µ*g of recombinant rat CCN2/CTGF (R&D Systems)(n=4) or 50 *µ*L of PBS as control (n=3) and subcutaneously injected the Matrigel mixture on the back of each mouse. At day 7 after injection, mice were euthanized. Matrigel plugs were removed, fixed in 4% paraformaldehyde in PBS, and embedded in paraffin. Sections (4 *µ*m thick) were deparaffinized and stained with hematoxylin-eosin (HE). Primary antibodies used for immunohistochemistry are listed in Table S3. The following secondary antibodies were used: Alexa594 goat anti-Syrian hamster IgG, Alexa488 donkey anti-rat IgG and Alexa488 donkey anti-rabbit IgG (Jackson Immunolaboratory, West Crove, PA, USA). Morphology was observed using a fluorescence microscope (BZ-710X; Keyence, Kyoto, Japan), and samples were examined using a confocal microscope (LSM710; Carl Zeiss, Tokyo, Japan).

### Statistical analysis

Statistical analysis was performed using GraphPad Prism software version 8. All experiments were performed more than three times, except the RTK assay (n=2), and data are expressed as the mean ± standard error (S.E.).

Comparative analysis was performed by Student’s t-test, one-way and repeated measures analysis of variance (ANOVA), or two-way ANOVA. For multiple comparisons, we performed Bonferroni or Sidak analysis. Statistical significance was set at a *p*-value of less than 0.05.

## Data availability

All data are contained within the article.

## Supporting information

This article contains supporting information.

## Acknowledgments

The authors thank Mrs. Yuriko Hamaguchi for assistance with animal experiments and histological analysis. We thank Gabrielle White Wolf, PhD, from Edanz (https://jp.edanz.com/ac) for editing a draft of this manuscript.

## Author contributions

S. H. and T. T. conducted and performed all experiments and wrote the manuscript. R.M. conducted preliminary experiments on CCN2. Se. K. and Sh.

K. supervised and arranged the experiments and reviewed and edited manuscript.

## Funding and additional information

This research was supported by a Grant-in-Aid for Scientific Research (grant no. 17K11866) from the Ministry of Education, Culture, Sports, Science and Technology of Japan; the Suisyo Kenkyu Project, Fukuoka University (grant no. 207202); and the Institute for Regenerative Medicine, Fukuoka University.

## Abbreviations

ERK: Extracellular signal-regulated kinase
IGF: Insulin-like growth factor
Tie: Tyrosine kinase with immunoglobulin-like and EGF-like domains
EGFR: Epidermal growth factor receptor
EphB4: Erythropoietin-producing hepatocellular receptor B4
ErbB4: Epidermal growth factor receptor 4
PDGFRα: Platelet-derived growth factor receptor alpha
MAPK: Mitogen-activated protein kinase
JNK: Jun N-terminal kinase
HSP: Heat shock protein
FGF: Fibroblast growth factors
TNFα: Tumor necrosis factor alpha
HUVEC: Human umbilical vein endothelial cell
ECM: Extracellular matrix
MSC: Mesenchymal stem cell
FBS: Fetal bovine serum
PBS: Phosphate buffered saline
BCA: Bicinchoninic acid
HRP: Horseradish peroxidase

## Supporting Information

Table S1. List of primers used for quantitative RT-PCR.

Table S2. List of siRNA sequences.

Table S3. List of antibodies used for Western blotting, immunoprecipitation assay and immunohistochemistry.

Figure S1. Phospho-RTK array.

Figure S2. Phosphorylation levels of p38 MAPK, JNK, AKT, NFκB, Smad2, 3, 1/5/9 in LECs treated with CCN2.

Figure S3. Itgb2 and Itgb3 expression levels in integrin β1-suppressed LECs.

Figure S4. CD31positive or β3-Tubulin positive tublar stracture was induced by CCN2 Matrigel plug.

**Table S1.**
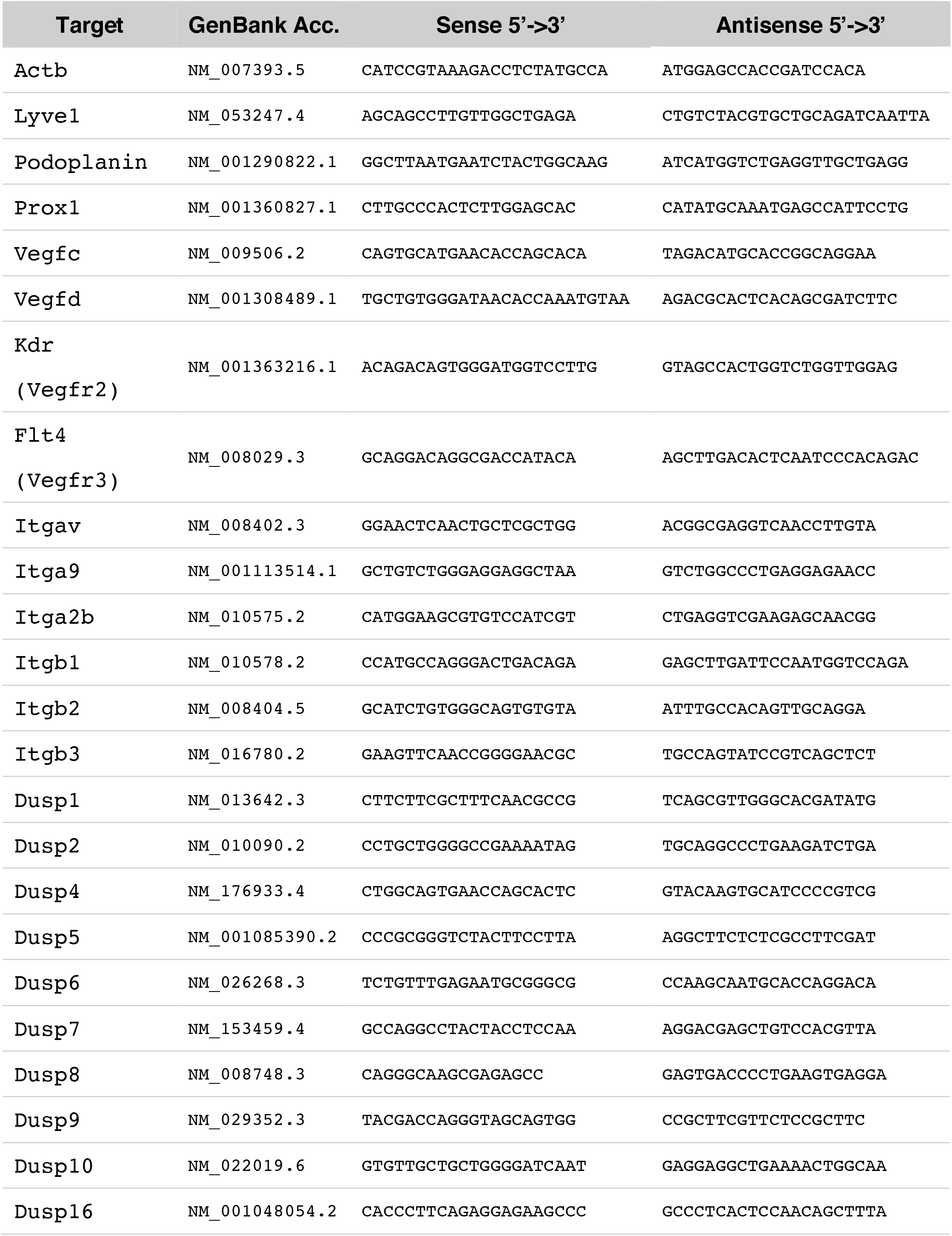
List of primers used for quantitative RT-PCR.

**Table S2.**
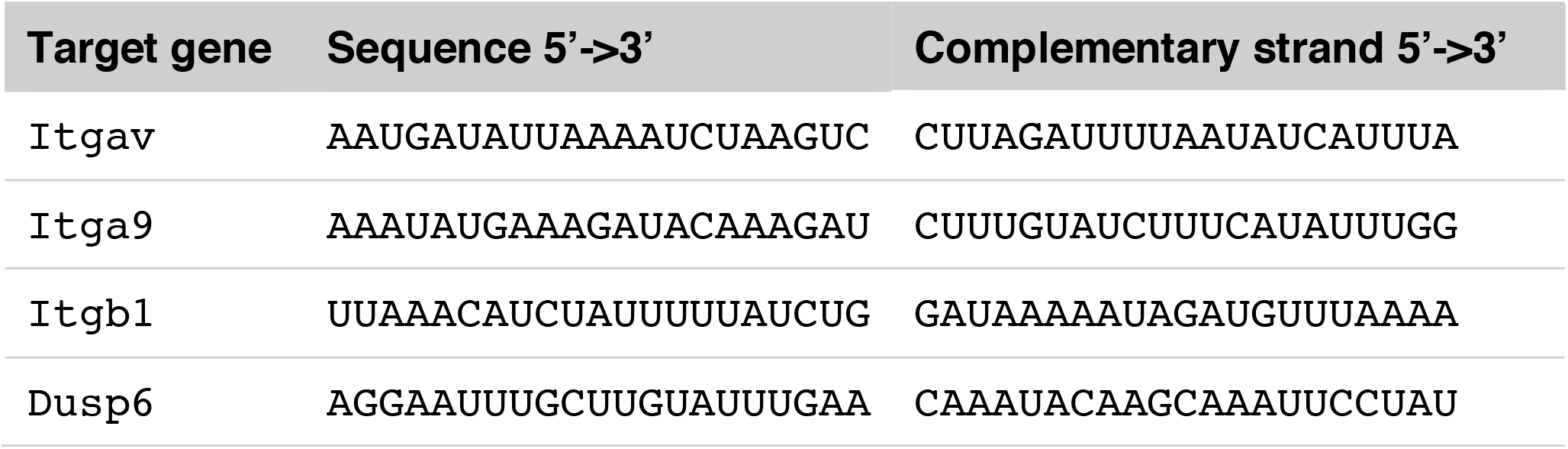
List of siRNA sequences.

**Table S3.** List of antibodies used for Western blotting, immunoprecipitation assay and immunohistochemistry.

| Target | Antibody | Supplier/<br>catalogue number |
| --- | --- | --- |
| phospho-ERK | Phospho-p44/42 MAPK (Erk1/2) (Thr202/Tyr204) Antibody | Cell Signaling Technology / 9101 |
| ERK (for WB) | p44/42 MAPK (Erk1/2) Antibody | Cell Signaling Technology / 9102 |
| ERK (for IP and IHC) | p44/42 MAPK (Erk1/2) (137F5) rabbit mAb | Cell Signaling Technology / 4695 |
| phospho-p38 | Phospho-p38 MAPK (Thr180/Tyr182) (3D7) rabbit mAb | Cell Signaling Technology / 9215 |
| p38 | p38 MAPK Antibody | Cell Signaling Technology / 9212 |
| phospho-JNK | Phospho-SAPK/JNK (Thr183/Tyr185) Antibody | Cell Signaling Technology / 9251 |
| JNK | SAPK/JNK Antibody | Cell Signaling Technology / 9252 |
| phospho-AKT | Phospho-Akt (Ser473) (D9E) XP rabbit mAb | Cell Signaling Technology / 4060 |
| AKT | Akt (pan) (C67E7) rabbit mAb | Cell Signaling Technology / 4691 |
| phospho-NFκB | Phospho-NF-κB p65 (Ser536) (93H1) rabbit mAb | Cell Signaling Technology / 3033 |
| NFκB | NF-κB p65 (D14E12) XP rabbit mAb | Cell Signaling Technology / 8242 |
| HSP90 | HSP90 Antibody | Cell Signaling Technology / 4874 |
| GAPDH | GAPDH (14C10) rabbit mAb | Cell Signaling Technology / 2118 |
| phospho-Smad1/5/9 | Phospho-Smad1 (Ser463/465)/<br>Smad5 (Ser463/465)/<br>Smad9 (Ser465/467) (D5B10) rabbit mAb | Cell Signaling Technology / 13820 |
| Smad1 | Smad1 (D59D7) XP rabbit mAb | Cell Signaling Technology / 6944 |
| Smad5 | Smad5 (D4G2) rabbit mAb | Cell Signaling Technology / 12534 |
| phospho-Smad2 | Phospho-Smad2 (Ser465/467) (138D4) rabbit mAb | Cell Signaling Technology / 3108 |
| Smad2 | Smad2 (D43B4) XP rabbit mAb | Cell Signaling Technology / 5339 |
| phospho-Smad3 | Phospho-Smad3 (Ser423/425) (C25A9) rabbit mAb | Cell Signaling Technology / 9520 |
| Smad3 | Smad3 (C67H9) rabbit mAb | Cell Signaling Technology / 9523 |
| Integrin αv | Anti-Integrin alpha V antibody [EPR16800] | abcam / ab179475 |
| Integrin α9 | Anti-Integrin alpha 9 antibody [EPR9722] | abcam / ab140599 |
| Integrin β1 | Anti-Integrin β1 Antibody, clone MB1.2 | R&S systems / MAB1997 |
| DUSP6 | Anti-DUSP6 antibody [EPR129Y] | abcam / ab76310 |

**Fig. S1.**
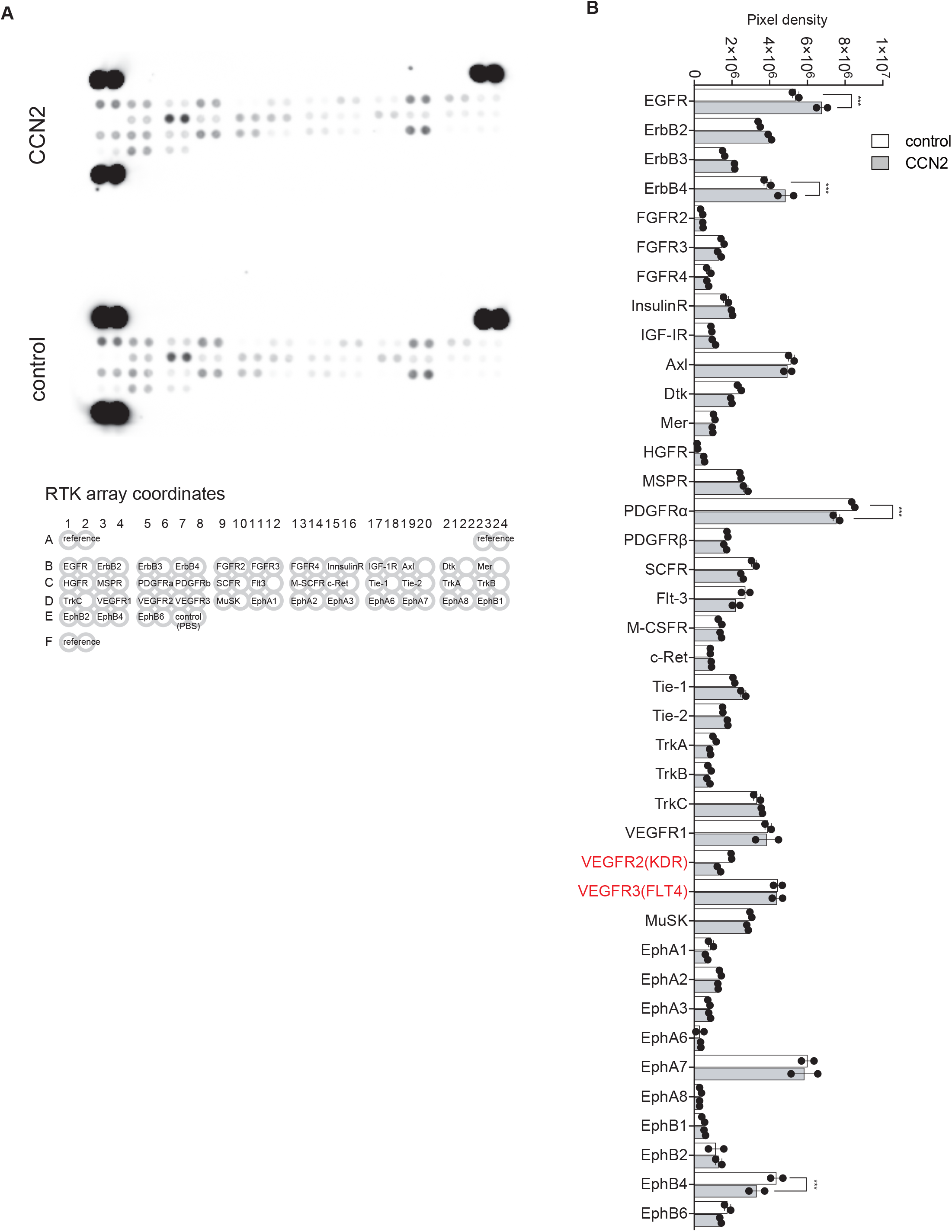
Phospho-RTK array. Cell lysate containing 200 *µ*g of protein prepared from LECs treated with 100 ng/mL CCN2 or PBS as (control) for 15 min was incubated with antibody-spotted membrane at 4°C overnight. Membrane was washed with wash buffer, incubated with anti-phospho-tyrosine antibody for 2hr. (A) Phosphorylated RTK was visualized with a image analyzer. (B) Pixel density of each RTK was measured by ImageQuant TL 8.1 software, and subtracted by that of reference. Two-way ANOVA, ^***^p < 0.001.

**Fig. S2.**
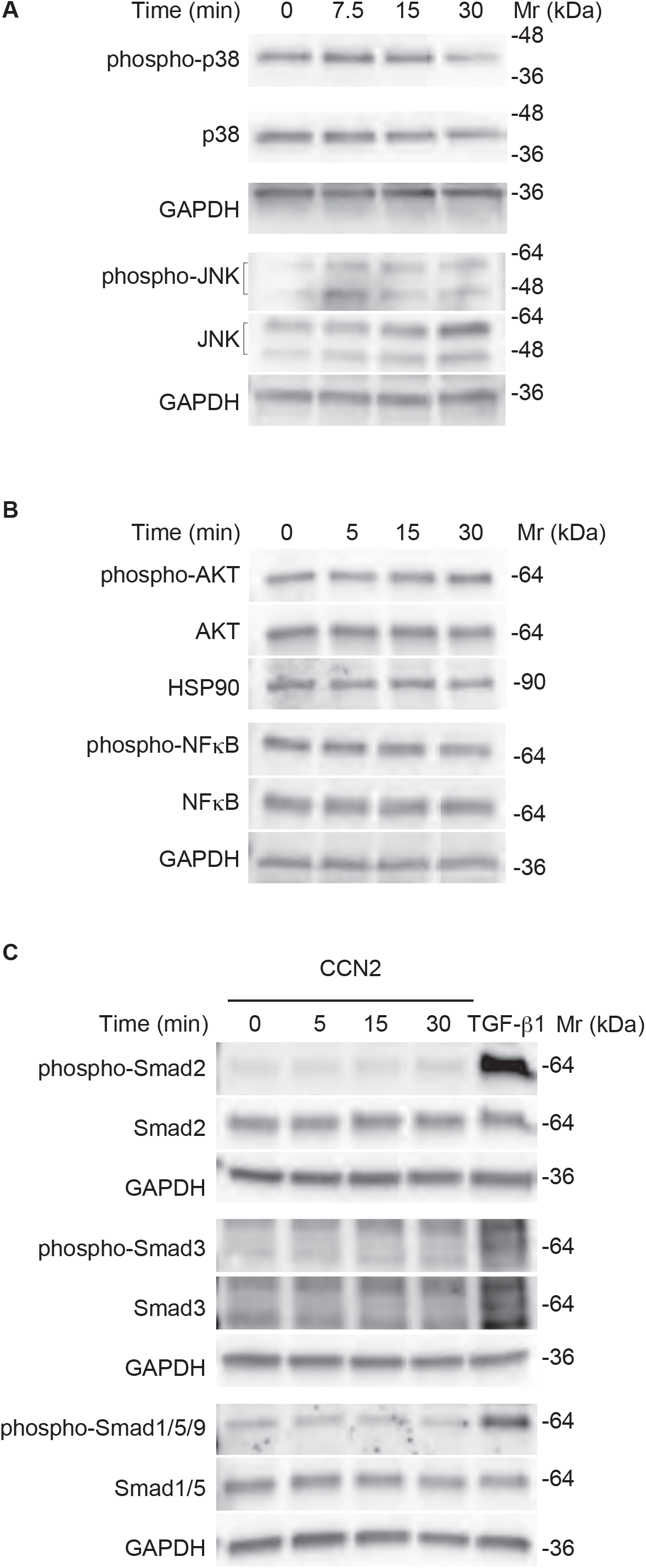
Phosphorylation levels of p38 MAPK, JNK, AKT, NFκB, Smad2, 3, 1/5/9 in LECs treated with CCN2. LECs were treated with 100 ng/mL of CCN2 for 0, 5 or 7.5, 10 and 30 min and lysed in RIPA buffer. 5-10 *µ*g of total protein was subjected to Western blotting. (A) phospho-p38 and phospho-JNK, (B) phospho-AKT and phospho-NFκB, (C) phospho-Smad2, phospho-Smad3 and phospho-Smad1/5/9. Cell lysate of LECs treated with TGFβ1 for 30 min was used for control for phospho-Smad3.

**Fig. S3.**
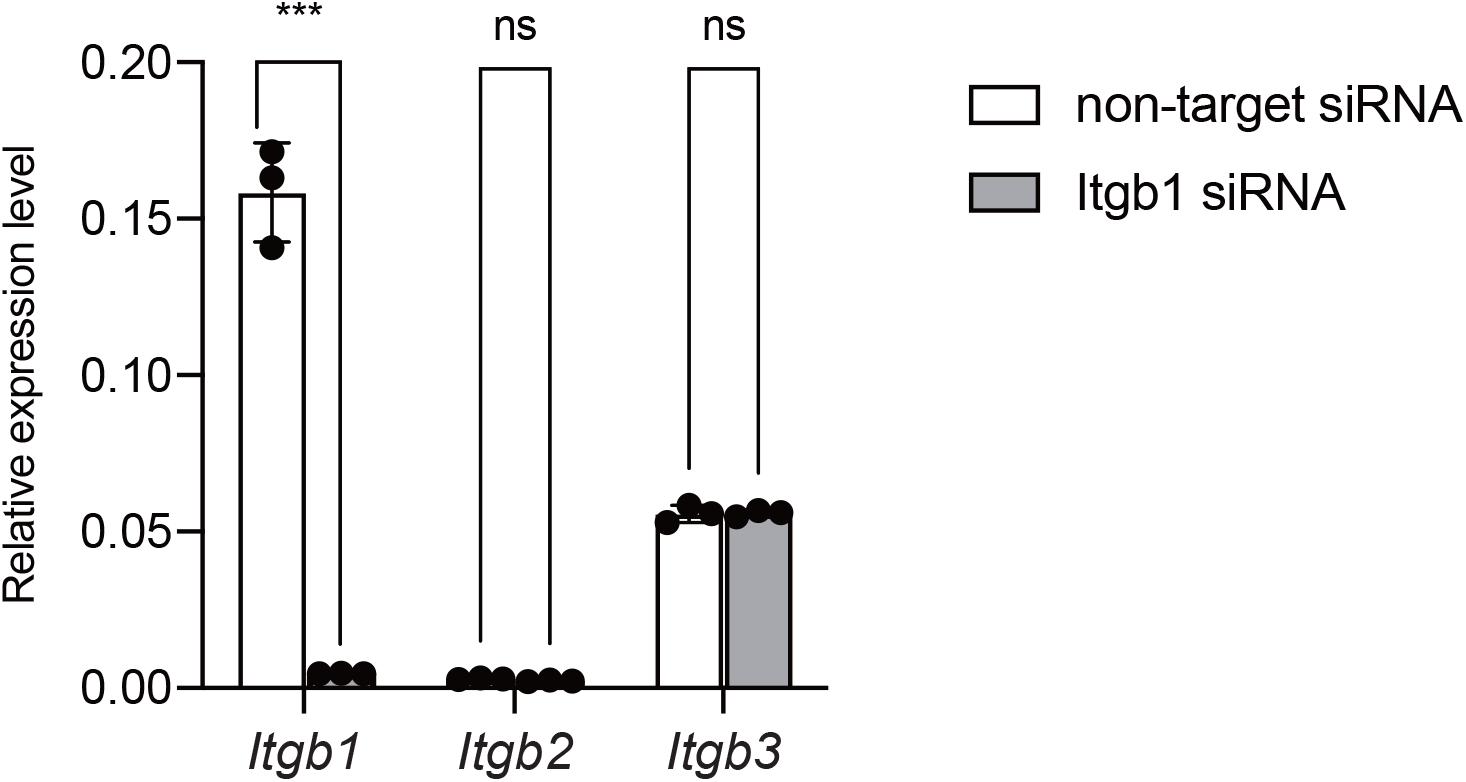
*Itgb2 and Itgb3 ex*pression levels in integrin β1-suppressed LECs. LECs were transfected with siRNA for Itgb1 or non-target siRNA, cultured for 24 hour, expression levels of *Itgb2* and *Itgb3* were analyzed with quantitative RT-PCR. Two-way ANOVA, ***p < 0.001. ns; not significant.

**Fig. S4.**
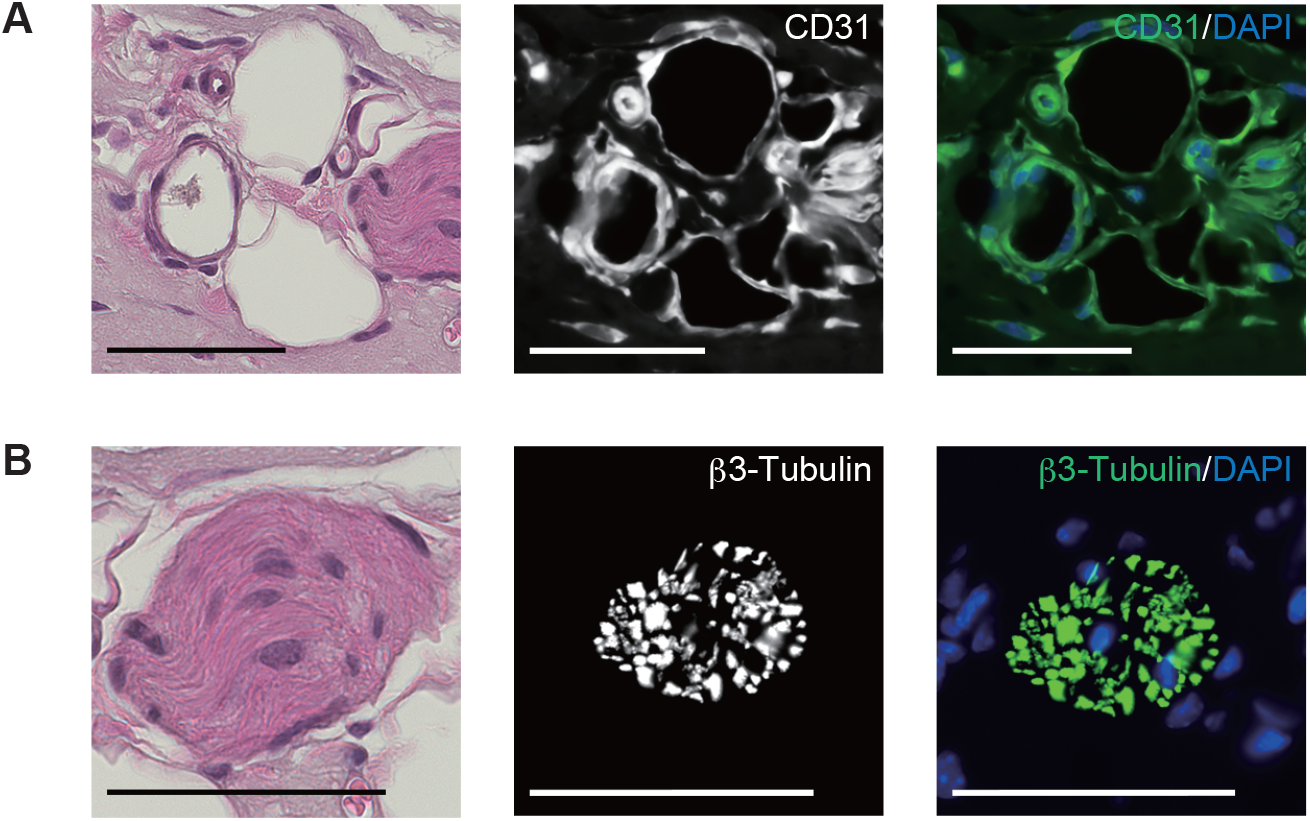
CD31 positive or β3-Tubulin positive tubular structure was induced by CCN2 in Matrigel plug. Matrigel and recombinant CCN2 were mixed and injected subcutaneously back of the mice. Control mice was injected with Matrigel and PBS mixture. Seven days after injection, Matrigel plug was removed, and expression of CD31 and β3-Tubulin was assessed by immunohistochemistry. (A) CD31 positive vessels were detected in CCN2 plug. (B) β3-Tubulin positive bundles detected in CCN2 plug. Scale bars: 50 *µ*m.

